# Is ChatGPT detrimental to innovation? A field experiment among university students

**DOI:** 10.1101/2024.04.03.588037

**Authors:** Mazen Hassan, Engi Amin, Sarah Mansour, Zeyad Kelani

**Author notes:** Corresponding author (SM). These authors contributed equally to this work.

## Abstract

ChatGPT represents a momentous technological breakthrough whose implications – along with other AI innovations – are yet to fully materialize. This paper is among the first attempts to experimentally test the effect of AI applications (in the form of ChatGPT) on three dependent variables usually assumed to be AI-collaterals: innovation, readiness to exert effort, and risk behaviour. We took advantage of the delayed introduction of ChatGPT in Egypt and conducted a pre-registered field experiment with nearly 100 senior university students at a public university. Over one month during term time, participants were asked to submit three graded essay assignments. In the treatment group, students were asked to write the essays using ChatGPT whereas in the control group, such option was neither mentioned nor allowed (the experiment was fielded before ChatGPT was legally operable in Egypt). One week after all assignments were submitted, the two groups were invited to the lab to play an innovation game (deploying multiple strategies to increase the sales of a hypothetical lemonade stand), a risk game (bomb risk elicitation task), and do a real effort task. The ChatGPT group was significantly less innovative, significantly less risk averse, and exerted less effort (however not statistically significant). Our results point to possible negative effects of AI applications but need further testing and larger samples to be confirmed.

## I. Introduction

Even before the rise of ChatGPT and other artificial intelligence text generators (AITGs), there has been a global debate – that is not short of controversy – on the risks and benefits of automation and artificial intelligence (1), with some calls for regulation (2), or at least a pause on developing the technology until further studies assess their impacts. The ascend of ChatGPT has certainly intensified this debate, with worries about the new technology from scientists, educators and even students (see for example (3)). Indeed, recent survey evidence has shown that two thirds of students use ChatGPT in their studies (4). In this context, this paper seeks to test the effect of the continuous usage of ChatGPT on innovation, effort levels, and risk-behaviour among university students.

Our assumption is straightforward and intriguing; that repeated reliance on technology to come up with answers to questions and to solve problems (just as ChatGPT does) creates a social norm of dependence on technology to innovate and – if used over a sustained period of time – crowds out the innovation drive by humans. To explore this question, we conducted a pre-registered field experiment with nearly 100 senior university students at a public university in Egypt where we tested the effect of using ChatGPT over a month in doing assignments on three dependent variables usually assumed by the literature to be AI-collaterals: innovation, risk behaviour, and readiness to exert effort. The field experiment extended for around four weeks and involved participants submitting three graded essay assignments during that period. In the treatment (ChatGPT) group, students were asked to write the essays using ChatGPT whereas in the control group, such option was neither mentioned nor allowed (the experiment was fielded before ChatGPT was legally operable in Egypt). One week after all assignments were submitted, the two groups were asked to take part in a lab experiment – without knowing what the treatment was – where they were asked to play an innovation and a risk game. Our findings are nuanced; the ChatGPT treatment group was significantly less innovative (measured by how frequently they changed the sales strategies at the 95% confidence level) and less risk averse (at the 90% confidence level). The treatment group also exerted less effort (measured by how frequently they recorded their strategies for their own reference over the rounds), although this result was not statistically significant.

To explain our theory and design in detail, from this point this paper is divided into five sections. The next section outlines our theory whereas section three presents the experimental design. Section four incudes the findings and section five concludes with a discussion of the possible limitations of the study and venues for future research.

## II. Theory

Technological advances have become a major factor affecting human behaviour and consequently the social norms generated from – and at a later stage, guiding – that behaviour. The invention of the personal computer and laptops has made working from home beyond office hours an expected behaviour from employers. The ascend of mobile phones has made people hardly unavailable. The rise of social media has affected the attention span of its users (5,6), given rise to social comparison and made us less social (7). This is hardly surprising given the penetration by technology of almost every aspect of our everyday life coupled by our increasing reliance on them to make our lives easier – and in parallel also different.

The premise of this paper is that technologies have spill-over effects across domains and are capable of affecting behaviour in areas and at times beyond those in which they are originally used. For example, it is by now an almost established fact that smart phones are making us less attentive in other tasks where we don’t use smart phones (8). They are also negatively affecting our recall accuracy and behavioural control (9,10), making their effect survive long after we stop using them – although of course such effects are dependent on duration and intensity of usage.

We contend that a similar spill-over effect happens with regard to one of the newest – but at the same time one of the fastest growing – technologies: ChatGPT. We argue that continuous usage of ChatGPT would also leave its imprints on socio-economic behaviour in domains where ChatGPT is not particularly used. We focus on three types of behaviour that are central to economic growth: innovation, effort and risk behaviour.

Starting with innovation, it is indeed a value so crucial to economic activity, productivity and even to social relations (e.g. 11,12). Innovation, however, is a mentally pressing task. It requires individuals to keep thinking about a certain task (and how to improve it) through a reiterative process of asking questions and trying to answer them. A central dimension of innovation therefore involves problem solving (13–15); whereby knowledge is produced via experimentation (16,17) and where early failure is an integral part of the process (18). ChatGPT – and other AITGs – however predominantly involve asking an algorithm a series of questions and waiting for the chatbot to come up with answers. If the task of answering the question is repeatedly outsourced however, over time the otherwise supposed innovator would not exert the same mental effort to answer the important questions and solve the pressing problems at hand, driving innovation (or at least human-driven innovation) down. Taking the mental exercise out of problem-solving is likely to lead to counter-innovation attitudes. Our first hypothesis therefore is:

H1: *The continuous use of ChatGPT over a sustained period of time, would drive down innovative behaviour*.

We do acknowledge that AI can also help innovation via different mechanisms. These might include freeing up the time of humans otherwise consumed in monotonous tasks (19) that are not structured to lead to innovation (although some studies do show that we might not be making use of this time in innovative tasks, but just in more screen time instead (20). Another mechanism could be AI helping in processing a large body of data to identify patterns that might unearth a problem and suggest its solution (see (14)). We argue however that these mechanisms are (a) more about incremental innovation (solving problem with a focus on one variable to be optimized), rather than the type of ground-breaking innovation that involves a radical shake-up of the status quo for the better, and (b) that they largely talk about machine-driven innovation (either partly or completely). In this paper however, we are interested in studying human-driven innovation and particularly whether the potential mechanisms outlined above would – over time – crowd out the innovation drive by humans that has been so central to human progress (21).

A second socio-economic attitude that could get affected by AITGs is risk behaviour. While a reasonable level of risk is required for economic activity to thrive, substantially low risk aversion could lead to reckless behaviour and endanger economic enterprises. We argue that the continuous usage of ChatGPT could increase risk tolerance via two mechanisms. First, there is the moral hazard context that this specific type of automation creates (22). The costless experimentation of asking a chatbot as many questions as one likes until one receives an answer (that one likes) is likely to prime users with a sense of insurance (23) that allows for endless experimentation at virtually no, or low cost, thereby decreasing their risk aversion. A second mechanism by which repeated usage of ChatGPT could affect risk behaviour is what previous research has shown that when humans interact with machines and algorithms, they tend to apply a different set of values: mainly getting less emotional and less concerned with social rules of conduct (for a review see (24)). Such decreased pro-sociality has been shown in trust games (25), in ultimatum games, as well as in dictator and public goods games (26). It is these social values however that many times restrict reckless behavior and make an individual stop short of taking major risks. Our second hypothesis therefore reads as follows:

H2: *The continuous use of ChatGPT over a sustained period of time, would decrease risk aversion.*

Effort is usually a likely collateral of many technological advances. Because most technologies mainly aim at making a machine, a software or an algorithm replace some aspect of human effort (e.g. sending voice message instead of typing it, voice or face recognition instead of pressing buttons), increased reliance on technologies are likely to decrease our perception of the amount of effort required from us, leading to complacency (27). Relevant research has shown how technology has made users less active (28). Such effects are not only confined to people with prior low skills: (29) have shown that the use of a spellcheck program makes even individuals with high writing skills miss spelling errors later on. In health care domains, it was also shown that the capacity of professional staff in reading mammograms went down (making them miss cancers) if they used computer-aided detection systems (30). Our third hypothesis therefore reads as follows:

H3: *The continuous use of ChatGPT over a sustained period of time, would decrease the level of effort one exerts to complete a task*.

## III. Methods

We conducted a pre-registered field experiment in which 94 senior students from an Egyptian public university participated. We chose a university as our experimental context for two reasons. First, universities and students are usually expected to be primary engines for innovation, by developing novel ideas, debating big questions, and having the time to experiment with multiple solutions. Indeed, most breakthroughs have usually originated from university campuses (31). Second, the potential negative effect of ChatGPT on university students was among the first worries raised when the technology made its debut. Just as artificial intelligence text generators (AITGs) started to make headlines, many feared that students would use the chatbot to do assignments, write theses, and hence change – or harm – their learning trajectory (for a discussion, see (32–35). This worry was deepened after several studies pointed to a reasonable performance by ChatGPT in handling academic tasks, such as taking exams and generating academic abstracts (36). A university campus, therefore, seemed to be an ideal venue for a field experiment on the probable effects of ChatGPT on innovation.

We designed our experiment to mimic educational institutions’ most basic concerns about ChatGPT: whether students seeking the assistance of the technology would be less innovative over time. Our participants were undergraduate students who had the same major (social sciences) and minor (social science computing) but were enrolled in two different courses: *Social Network Analysis* and *Data Mining*. We used the enrollment in either of these two courses as our unit of randomization to assign participants to either the treatment or the control. We took this decision to avoid potential across-treatment contamination if the intervention was applied within the same course. In the next section, we present analyses showing that both groups (the treatment and the control) were not significantly different in crucial background variables. The field component of the experiment was administered as follows. Students in both courses (i.e. the treatment and the control) were asked to submit three graded essay assignments over one month during term time in April 2023. The difference between both groups was that members of the treatment group (enrolled in the *Social Network Analysis* course) were asked to write the essays using ChatGPT, whereas for members of the control group (enrolled in the *Data Mining* course), ChatGPT option was neither mentioned nor allowed.

The time in which we conducted the experiment enabled us to use a small window of opportunity when access to ChatGPT was still unavailable in Egypt. It was only in November 2023 that Egyptians could create an account and access ChatGPT with local cell phone numbers and Egyptian IP addresses. While before November 2023 Egyptians could access ChatGPT via a VPN and foreign phone number, this step was largely burdensome and hence was quite rare at the time. For the treatment group, we created ChatGPT accounts for them, using foreign cell phone numbers. This context allowed for a field experiment that minimizes the risk of contamination – where subject in the control group might also use ChatGPT – and hence ensured more reliable results.

One week after all assignments were submitted, the two groups were asked to take part in a lab experiment. At the beginning of each session, the experimenter would greet the participants, give them general rules, and ask if they would like to participate. Anyone who did not want to participate was free to leave. A screen with a consent form was prompted before participants were allowed to start the experiment. Only participants who consented were allowed to proceed with the experimental instructions. If a participant did not explicitly click on the consent button, the experiment was terminated for them immediately. A screenshot of the consent form presented before the commencement of the actual experiment can be found as part of the experimental script in the supporting documents. To have a signed documentation from the participants, after the experiment had concluded and the payoffs were shown, a receipt slip was given to each participant to fill in and sign with the amount they received. This ensured that participants did not provide any identification during the experiment or on the computers to protect their anonymity and that the experimenters had official signed documentation of participation (so no participant would be allowed to enter the experiment again).

In the lab experiment subjects were asked to play an innovation game (37) and a risk game (38). In the innovation game, participants had to innovate – over seven rounds – to increase the sales of a hypothetical lemonade stand by deciding on the values of six parameters (after having received advice from the previous manager):

a. location of the stand (school, business, stadium).
b. lemonade color (green, pink).
c. sugar content (scale from 0 to 10).
d. lemon concentration (scale from 0 to 10).
e. price (scale from 0 to 10).
f. Writing an advertising message or slogan with a maximum of 20 words.

After each round, participants were shown the profit they made using the parameter values they had chosen and were asked whether they wanted to change or keep their parameter choices for the next round. Payoffs were calculated according to the optimal parameters set by (37) in which any deviation from the optimal parameter values (which were unknown to the subjects) was penalized. We added to the design by (37) a further innovation task that asked participants to write an advertising message or slogan. This additional task was not included in the payoffs structure but was used to further assess innovation as will be shown below. After the conclusion of the sessions, four external annotators were tasked with rating the advertising messages on a creativity scale from 0 to 10. The annotators were unaware of the research question or the experimental design to ensure coding impartiality.

To measure effort levels, at the beginning of each session, participants were given a sheet of outlined paper to record their strategies for reference over the rounds. Recording previous choices should have been essential for supposedly efficient participants in order to avoid repeating their choices in future rounds (given the many decisions they had to make in each round). Filling in the cells was voluntary, and participants were not told that recording their decisions and profits would be measured. Instead, they were told that the sheet was for them *if* they wanted to track their choices and profits. These sheets were collected after the session concluded. We used the number of cells filled out in the sheets as our measure of participants’ effort levels.

To measure risk behaviour, participants completed a dynamic version of (38) “Bomb Risk Elicitation Task.” The task is a game that presents participants with 100 closed “boxes,” 99 of which contain a small monetary reward (the amount in each of these 99 boxes is the same). However, one of the 100 boxes contains a bomb that if opened would wipe out all monetary gains achieved thus far. Once the game starts, a box gets collected every second until the participant clicks “stop” or all 100 boxes are collected. The underlying assumption is that risk-tolerant individuals will collect more boxes while risk-averse individuals will collect fewer. See S1 for the experimental script.

## IV. Results

Table 1 shows a summary of the characteristics of the participants in the control and treatment groups. A total of 94 subjects were initially recruited for the experiment. However, one subject was discarded due to missing information during payouts. The majority of the participants were female – which is common in social science departments in Egypt (e.g. (39–41)). Average age was 21 and around 92% were Muslims. Balance tests conducted to compare the demographic composition of the control and treatment group revealed no randomization failures (t-test for age and financial status, chi-square tests for gender, religion, and residence; failed to reject Ho, p-value > 0.05).

**Table 1.**
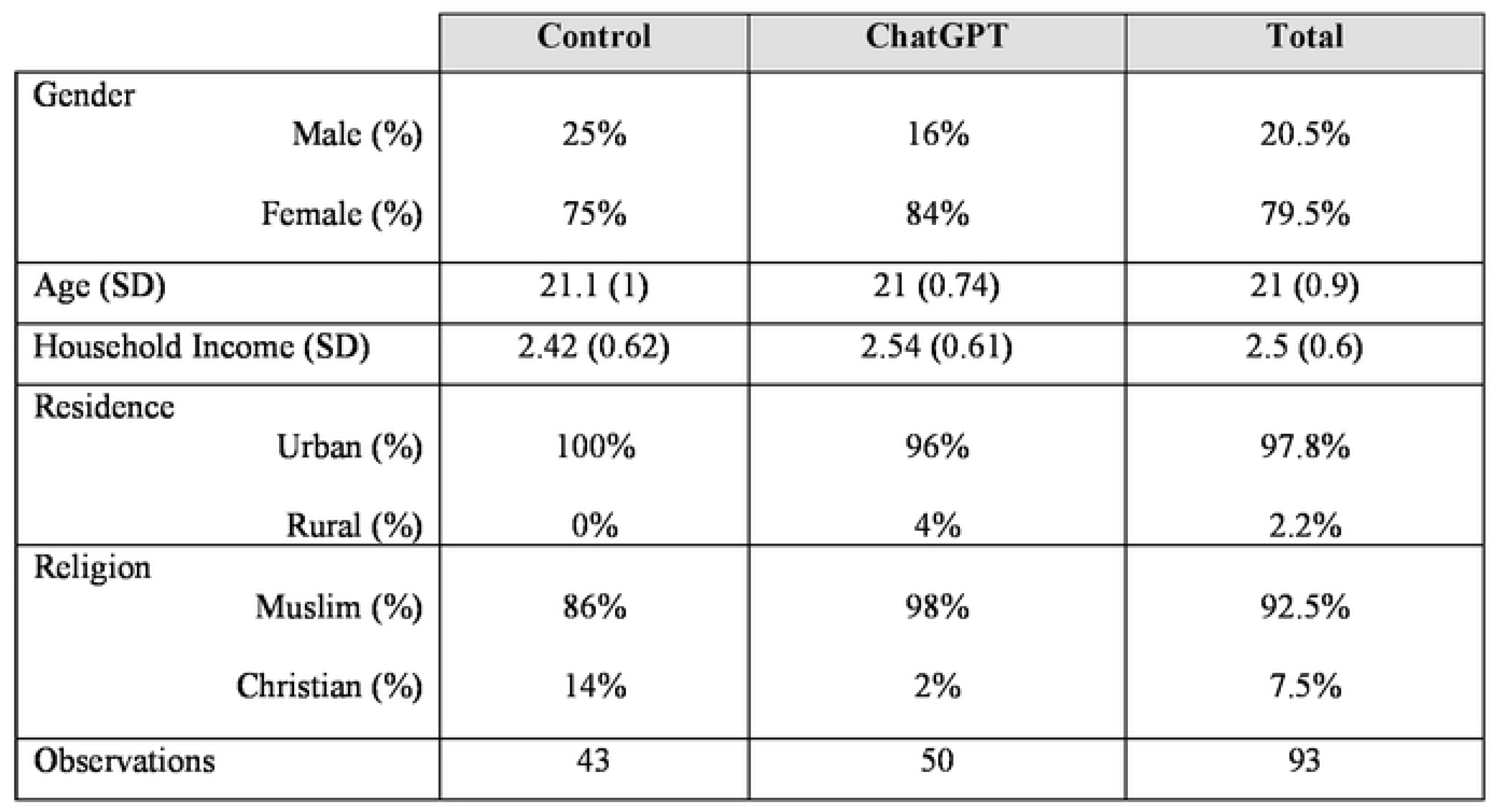
Subject Demographics.

In the following, we investigate whether the use of ChatGPT for four weeks before the lab experiment influenced (i) the innovative behaviour based on the lemonade stand game, (ii) the level of effort calculated from the sheets used to record strategies, and (iii) the risk behaviour concluded from the bomb risk elicitation task. Data across sessions was compiled and cleaned. After performing initial outlier analysis, two anomalous observations were detected in the dataset, and we decided to remove them based on interquartile range method (1.5xIQR rule).

### First: Innovative Behaviour based on the Lemonade Stand Game

To measure the level of innovativeness from the lemonade stand game, we started by looking at whether there is significant treatment effect on any of the three continuous decision parameters (sugar content, lemon concentration, and price) that participants had to decide on in each round. Since we measured innovative behavior by the subject-specific standard deviation, a la Ederer and Manso (37), we focused only on the three continuous decision parameters (sugar content, lemon concentration, and price). Although this is a random first check – because theoretically we do not expect one parameter to be more conducive to innovation than the rest – it still is an important first step to look at the parameters separately. Fig 1 below shows that a significant difference in innovative behaviour (measured by the subject-specific standard deviation) exists in the decision regarding lemon concentration (Mann Whitney p-value=0.036). There is no significant difference however with regard to the sugar content or the price when comparing the treatment to the control (Mann Whitney p-value > 0.1).

**Fig 1.**
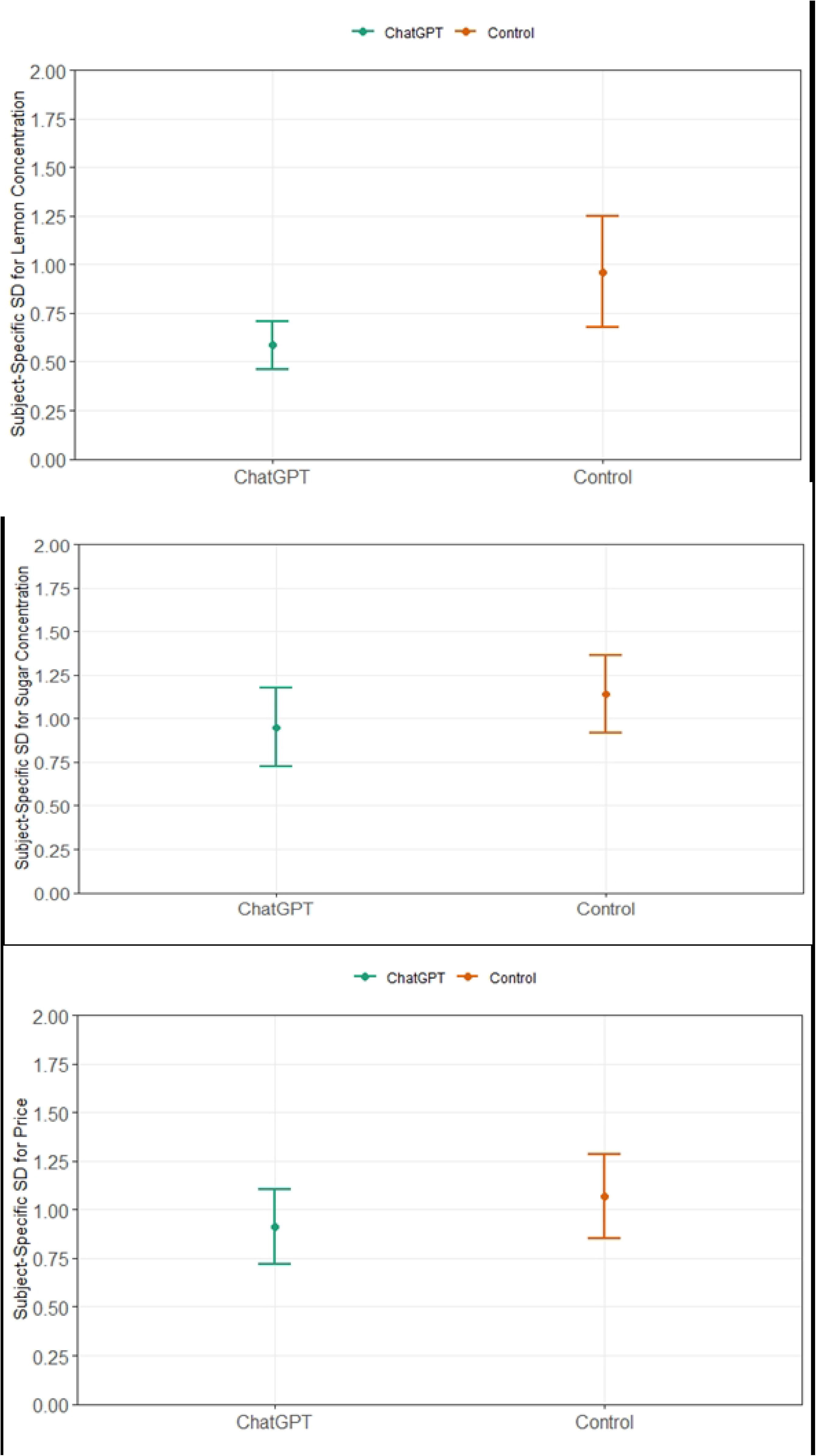
Average Level of Innovativeness for Control and ChatGPT Groups (Individual parameters)

Next, we move to measure innovativeness by examining the propensity of the subject to explore different values of the three parameters combined (sugar content, lemon concentration, and price), thus capturing the degree of novelty in the sales strategy. This was calculated as the average, subject-specific standard deviation of strategy choices for the three continuous variables (sugar content, lemon content, and price). Fig 2 shows the average level of innovativeness among the control and ChatGPT treatment groups. The control group had a significantly higher level of innovativeness when compared to the ChatGPT group (Mann Whitney U test, p-value = 0.037), confirming H1.

**Fig 2.**
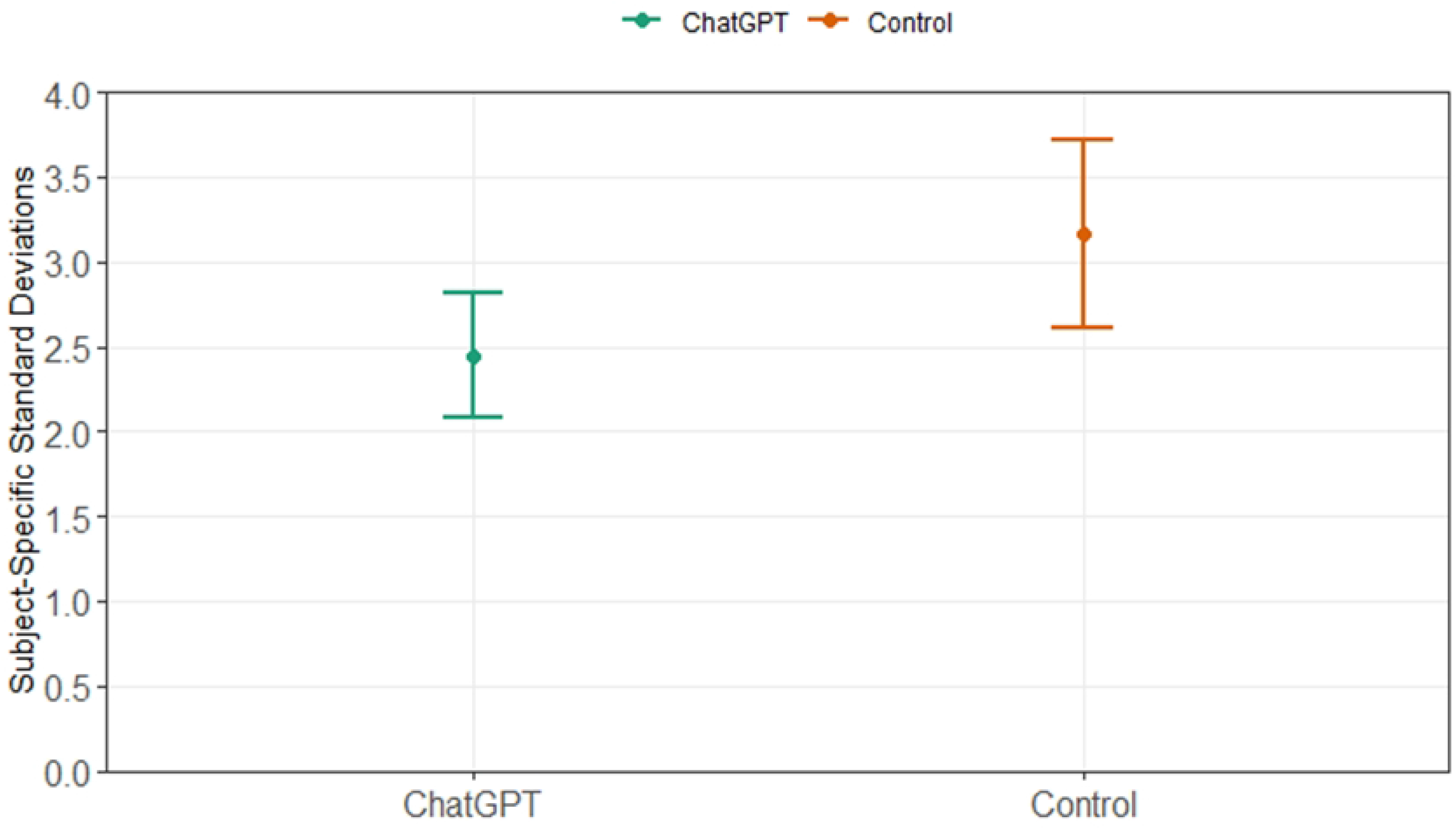
Average Aggregate Level of Innovativeness for Control and ChatGPT Groups.

Table 2 presents the results of regression analyses on two models with the level of innovativeness as the dependent variable, after controlling for potential confounders. The coefficient for ChatGPT is negative and statistically significant across the two models (at the 95% confidence level in model 1 and at the 90% level in model 2), indicating that the level of innovation for participants in the ChatGPT treatment is *significantly lower* than the control group, after controlling for other variables. The negative values suggest that ChatGPT usage is associated with a decrease in innovation scores by approximately 0.6 to 0.72 standard deviation points, depending on the model. Additionally, living in an urban area seems to have a positive and significant effect on innovation in the second model. Moreover, household income is negatively associated with innovation and is statistically significant, but only at the 0.1 level, suggesting a moderate confidence level that having higher income may reduce innovation scores. This could be because participants on higher incomes might not have valued the need to innovate to increase their payoffs as those on lower incomes.

**Table 2.** Regression Analysis for Level of Innovativeness in Lemonade Stand Game.

|  | *Dependent variable:* |  |
| --- | --- | --- |
|  | Exploration of sales strategies |  |
|  | (1) | (2) |
| ChatGPT | -0.719**<br>(0.326) | -0.604*<br>(0.328) |
| Female |  | -0.320<br>(0.462) |
| Age |  | -0.030<br>(0.173) |
| Urban |  | 0.673**<br>(0.322) |
| Financials |  | -0.510*<br>(0.273) |
| Constant | 3.167***<br>(0.271) | 4.081<br>(3.529) |
| Observations | 91 | 91 |

*Notes: Results are clustered by Subject ID, \*\*\* p<0.01, \*\* p<0.05, \* p<0.1*

The second innovation test we ran was coding of the advertising messages that participants were asked to write in each round along an innovation scale from 1 to 10. This task culminated in 265 unique messages (as some participants chose not to change their messages at some rounds). The coding was done by four coders after the experiment. Fig 3 below shows no significant difference in the creativity of these messages between the treatment and the control (Mann Whitney U test, p-value > 0.05), running against H1. We also ran other tests tracing innovation that however produced insignificant results. These can be found in appendix II.

**Fig 3.**
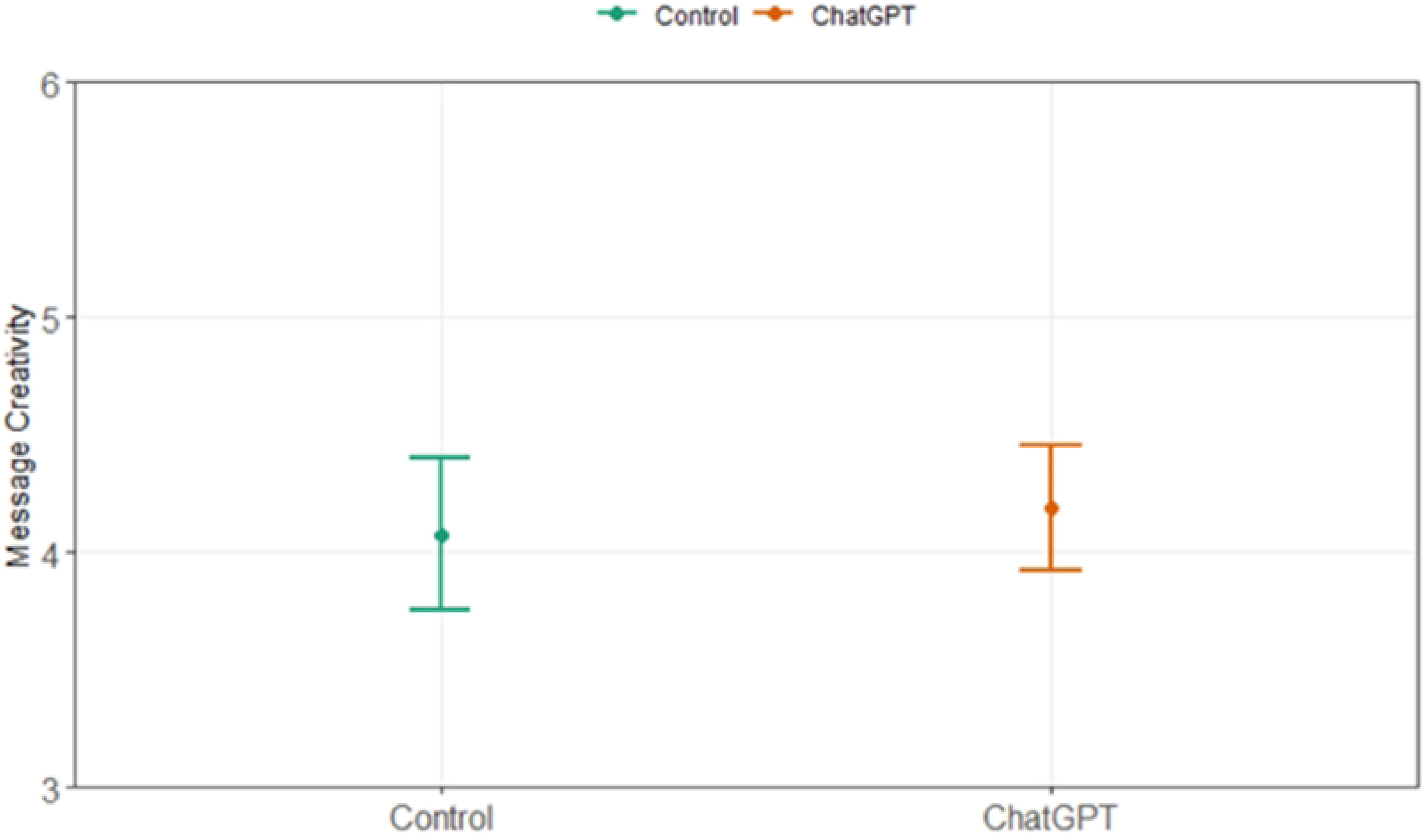
Creativity scores of advertising messages between Control and ChatGPT Groups.

### Second: Risk Behaviour from the Bomb Risk Elicitation Task

To calculate risk behaviours from the Bomb Risk Elicitation Task, the propensity for risk-taking among participants was calculated using the number of boxes they left unopened before they decided to press the *stop* button. A larger number of boxes opened reflects a higher inclination towards risk-taking, as the probability of encountering a bomb that would destroy all earnings increases. A subject was considered risk-loving if s/he opened more than 1/3 of boxes and risk averse otherwise. Fig 4 illustrates the proportion of risk lovers in both the control group and the ChatGPT treatment. The proportion of risk lovers in the ChatGPT group was higher than that of the control group, as suggested by H2. Moreover, this observed difference is statistically significant (Mann Whitney U test, p-value = 0.079).

**Fig 4.**
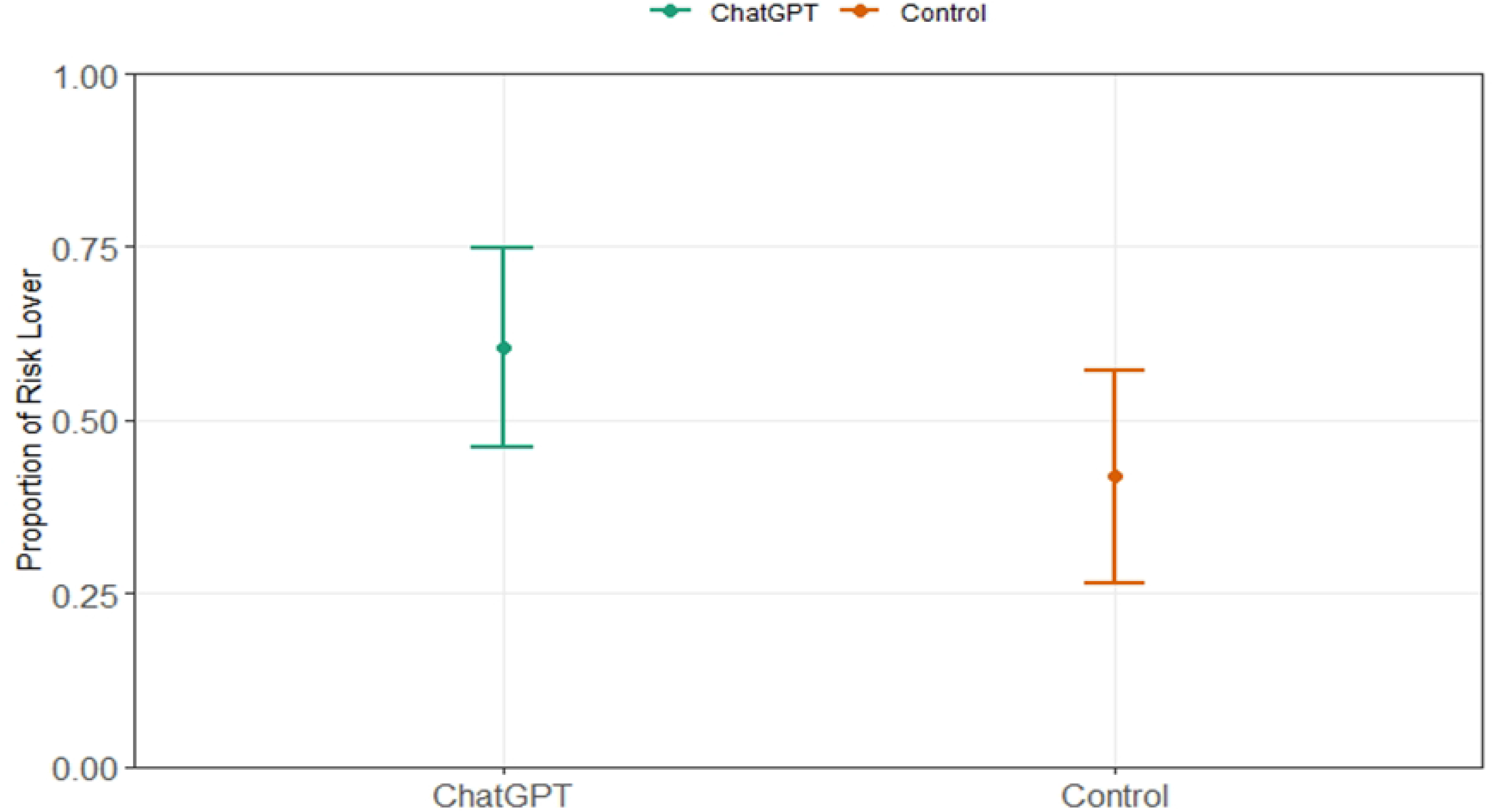
Proportion of Risk Lovers among Control and ChatGPT Groups.

Table 3 presents regression analyses investigating the likelihood of risk loving behaviour. Across both models, the coefficients for the ChatGPT treatment are positive (at the 90% confidence level), indicating that individuals in the ChatGPT treatment group *are more likely* to become risk lovers than those in the control group (which is in line with the hypothesis). Moreover, these results are statistically significant. In the second model, being a female is negative and statistically significant, indicating a lower likelihood of being a risk loving person.

**Table 3.** Regression Analysis for Risk Loving Behavior in the Bomb Risk Elicitation Task.

|  | *Dependent variable:* |  |
| --- | --- | --- |
|  | Risk Lover |  |
|  | (1) | (2) |
| ChatGPT | 0.751* | 0.818* |
|  | (0.427) | (0.449) |
| Female |  | -1.135* |
|  |  | (0.631) |
| Age |  | -0.002 |
|  |  | (0.252) |
| Urban |  | 0.236 |
|  |  | (1.484) |
| Household Income |  | 0.639 |
|  |  | (0.400) |
| Constant | -0.329 | -0.608 |
|  | (0.309) | (5.386) |
| Observations | 91 | 91 |
| Akaike Inf. Crit. | 126.909 | 130.316 |
| *Note:* *** p<0.01, ** p<0.05, * p<0.1 |  |  |

### Third: Level of Effort as Elicited from the Sheet for Recording Strategies

To assess the level of effort participants exerted (measured by their recording of the sales strategies employed at each of the seven rounds), we calculated the number of cells filled out by each subject at each period (min=0 and max=8). As mentioned above, filling in the cells was voluntary, and participants were not told that recording their decisions and profits would be measured. Fig 5 shows the sheet given to subjects at the beginning of the session if they wanted to record their choices and profits over the rounds.

**Fig 5.**
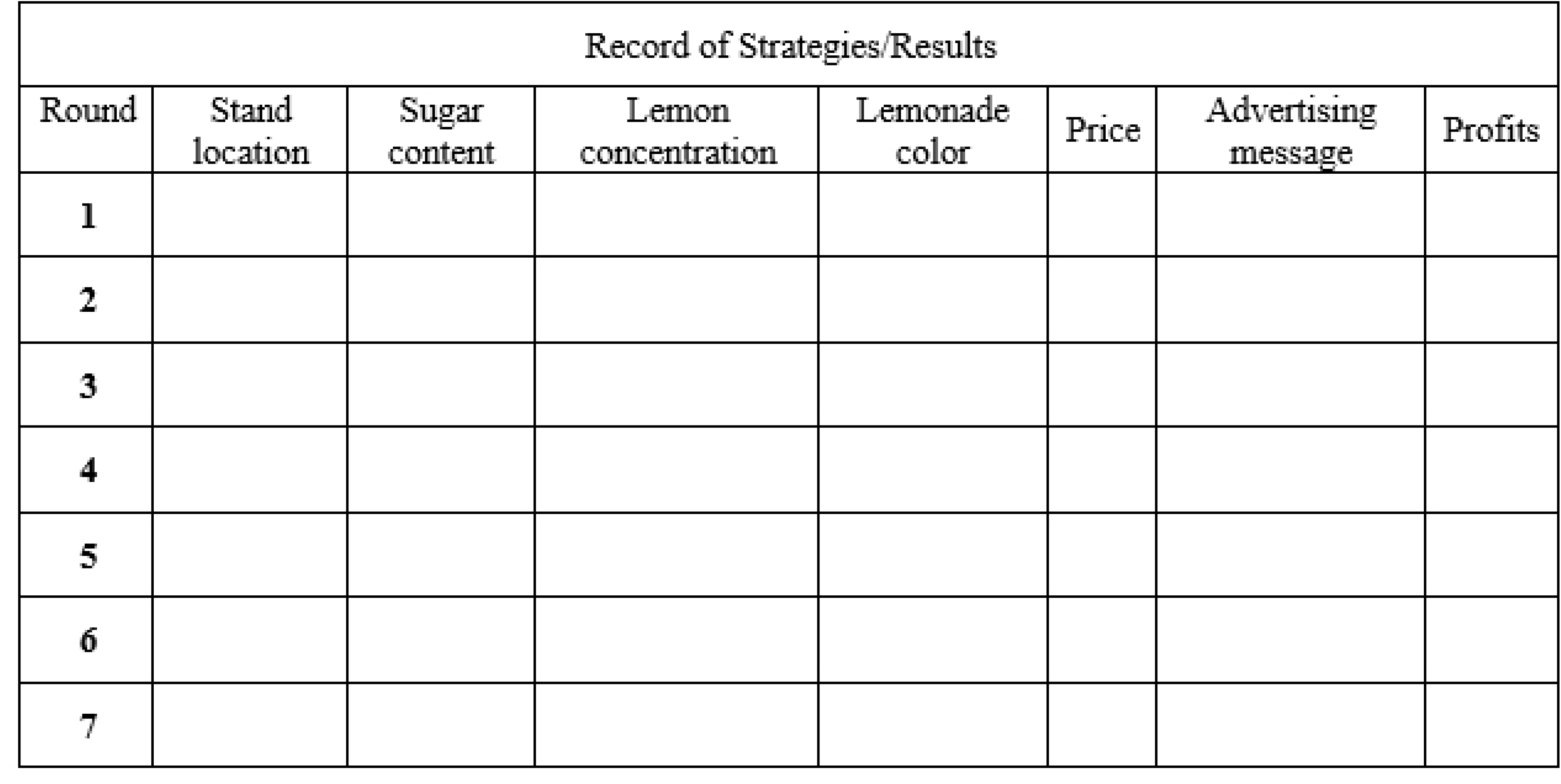
Sheet for Eliciting Effort.

Fig 6 displays the average effort exerted by the control and ChatGPT groups. It shows that the average effort exerted by participants in the control group is slightly higher than that exerted by the ChatGPT treatment group which runs in line with H3. However, this difference is statistically insignificant (Mann Whitney U test, p-value > 0.05).

**Fig 6.**
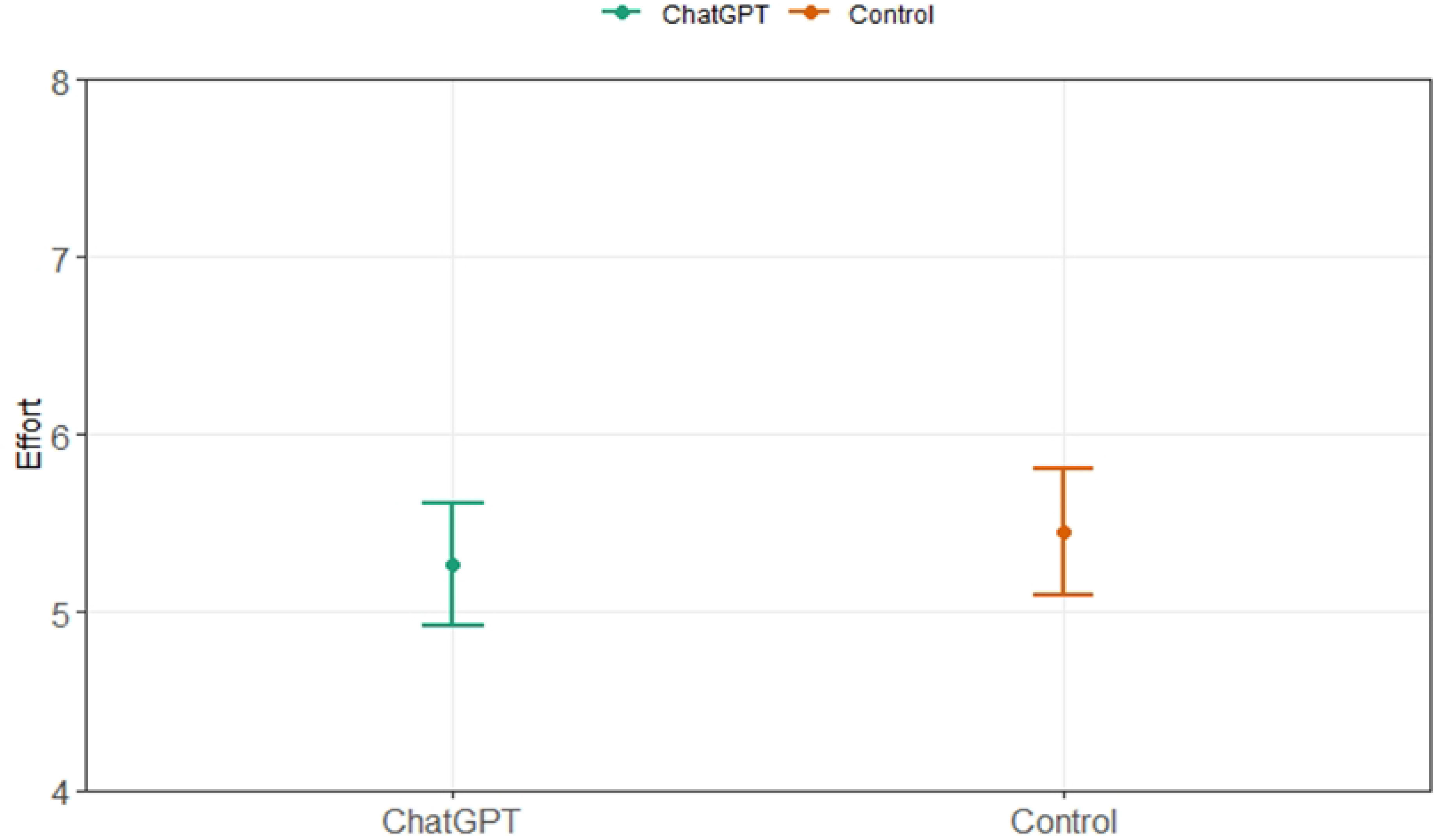
Average Level of Effort for Control and ChatGPT Groups.

Table 4 presents the results of regression analyses on three models with the level of effort as the dependent variable. Across all models, the coefficient for the ChatGPT treatment is negative (as expected by the hypothesis) suggesting that participants using ChatGPT exerted slightly less effort than the control. However, these results are not statistically significant in any of the models. The coefficient for the period is negative and is statistically significant in models 2 and 3 at the 0.1 significance level, implying that as the game progressed, participants tended to exert less effort, possibly due to fatigue or diminishing interest. In model 3, residing in an urban area is associated with a significantly higher level of effort, suggesting that urban participants were more engaged or able to exert more effort in the game.

**Table 4.** Regression Analysis for Level of Effort Exerted in Lemonade Stand Game.

|  | *Dependent variable:* |  |  |
| --- | --- | --- | --- |
|  | (1) | Effort<br>(2) | (3) |
| ChatGPT | -0.184<br>(0.615) | -0.184<br>(0.615) | -0.035<br>(0.611) |
| Female |  |  | -0.206<br>(0.852) |
| Age |  |  | -0.411<br>(0.392) |
| Urban |  |  | 4.285***<br>(0.681) |
| Household Income |  |  | -0.187<br>(0.546) |
| Period |  | -0.216***<br>(0.041) | -0.216***<br>(0.041) |
| Constant | 5.449***<br>(0.449) | 6.313***<br>(0.432) | 11.106<br>(7.963) |
| Observations | 637 | 637 | 637 |

*Notes: Results are clustered by Subject ID, \*\*\* p<0.01, \*\* p<0.05, \* p<0.1*

Putting together the details of the full picture, our results seem to be nuanced. Exposure to ChatGPT over a few weeks in doing assignments was shown to significantly decrease innovation, as measured by our lemonade stand game. Moreover, the use of ChatGPT positively affected risk-taking behavior compared to the control, and the effect was statistically significant. On the other hand, ChatGPT also had a negative impact on the level of effort exerted in a task, albeit not statistically significant. The fact that one of our two statistically significant findings is at the 90% confidence level however highlight the need for further testing.

## V. Discussion

The fast spread of ChatGPT has changed the nature of human-computer interaction, providing users with sophisticated conversational interfaces that respond intelligently to different prompts and queries. In this paper, we examined the effect of continuous reliance on such a technological breakthrough when doing assignments on undergraduate students’ innovative behaviour, risk attitudes and readiness to exert effort. Our findings have shown significant results on innovation and risk, but not on effort levels.

We would like to use this discussion section to talk about some caveats and possible limitations of our study, as well as venues for future research. On caveats, despite the focus of the paper and its pre-registered hypothesis on possible negative effects of ChatGPT, it is still important to highlight that AI applications and AITGs are also likely to have significant advantages. Taking the social clues out of the decision making, for example, was shown to produce less discriminatory decisions than human decisions (e.g. (42,43)). AI also has a huge potential in dealing with a size of data that is simply too large for humans to handle effectively. Depending on usage techniques, even ChatGPT itself could contribute to the development of innovation, if it succeeds – through its provision of a vast amount of valuable information – to foster a culture of curiosity and continuous exploration (see (44)) among its users. This study therefore does not deny the multiple advantages that AI and AITGs can generate.

On limitations, the findings of our study should be read in light of possible design restrictions. Firstly, there is the relatively small sample size on which we were restricted by the number of students already registered in the two courses we had access to. Secondly, the worry about spill-over effect between treatment and control that made us take course registration as the criterion for randomization which – ideally – is not the best randomization strategy. Further testing of our preliminary findings whilst improving on these two possible limitations therefore is important.

Finally, on venues for future research, although field experiments will always have the advantage of high ecological validity, testing the effect of AITGs on behaviour could also benefit from the controlled environment of lab experiments. In such designs, spill-over worries would be minimal. It is also important to note that we only tracked one dimension of innovation; that is innovation related to problem solving. There are however many other dimensions of innovation. One such dimension, for example, is the active reflection that individuals engage with occasionally (14). Such aspect of creativity and innovation requires a free mindset; one that does not specifically think in terms of a specific problem but that wanders around, accumulates ideas that could be used at later times (45).

## Acknowledgments

The authors would like to thank Dr. Heba Medhat Zaki and Dr. Mayada Aref for giving us access to their courses to apply both the field and lab interventions on their students. The authors would also like to thank the IT team at the experimental lab of the Faculty of Economics and Political Science, Cairo University and Nourhan Abdelhamid Elsheikh for their dedication throughout the experimental sessions.

## List of References

1. Pasquale F. The black box society: The secret algorithms that control money and information. Harvard University Press; 2015.

2. Wachter S, Mittelstadt B. A right to reasonable inferences: re-thinking data protection law in the age of big data and AI. Columbia Bus Law Rev. 2019;494.

3. Farhi F, Jeljeli R, Aburezeq I, Dweikat FF, Al-shami SA, Slamene R. Analyzing the students’ views, concerns, and perceived ethics about chat GPT usage. Comput Educ Artif Intell. 2023;100180.

4. von Garrel J, Mayer J. Artificial Intelligence in studies—use of ChatGPT and AI-based tools among students in Germany. Humanit Soc Sci Commun [Internet]. 2023;10(1):799. Available from: 10.1057/s41599-023-02304-7

5. Ophir E, Nass C, Wagner AD. Cognitive control in media multitaskers. Proc Natl Acad Sci U S A. 2009 Sep;106(37):15583–7.

6. Firth JA, Torous J, Firth J. Exploring the Impact of Internet Use on Memory and Attention Processes. Int J Environ Res Public Health. 2020 Dec;17(24).

7. Fischetti M. Social Technologies Are Making Us Less Social. Scientific American [Internet]. 2016 Oct; Available from: https://www.scientificamerican.com/article/social-technologies-are-making-us-less-social/

8. Altmann EM, Trafton JG, Hambrick DZ. Momentary interruptions can derail the train of thought. J Exp Psychol Gen. 2014 Feb;143(1):215–26.

9. Tanil CT, Yong MH. Mobile phones: The effect of its presence on learning and memory. PLoS One [Internet]. 2020 Aug 13;15(8):e0219233. Available from: 10.1371/journal.pone.0219233

10. Chen J, Liang Y, Mai C, Zhong X, Qu C. General Deficit in Inhibitory Control of Excessive Smartphone Users: Evidence from an Event-Related Potential Study. Front Psychol. 2016;7:511.

11. Amabile TM, Barsade SG, Mueller JS, Staw BM. Affect and Creativity at Work. Adm Sci Q [Internet]. 2005 Sep 1;50(3):367–403. Available from: 10.2189/asqu.2005.50.3.367

12. Dahl DW, Moreau P. The Influence and Value of Analogical Thinking during New Product Ideation. J Mark Res [Internet]. 2002 Feb 1;39(1):47–60. Available from: 10.1509/jmkr.39.1.47.18930

13. Amabile TM. Creativity, Artificial Intelligence, and a World of Surprises. Acad Manag Discov [Internet]. 2019 Feb 27;6(3):351–4. Available from: 10.5465/amd.2019.0075

14. Bieser J. Creative through AI – How artificial intelligence can support the development of new ideas [Internet]. Gottlieb Duttweiler Institute Research Paper No. Forthcoming. 2022. Available from: https://gdi.ch/en/publications/studies/creative-through-ai-pdf-2022-e#attr=

15. Sternberg RJ, Lubart TI. The concept of creativity: Prospects and paradigms. Handb Creat. 1999;1(3–15).

16. Arrow KJ. Classificatory Notes on the Production and Transmission of Technological Knowledge. Am Econ Rev [Internet]. 1969 Mar 26;59(2):29–35. Available from: http://www.jstor.org/stable/1823650

17. Weitzman M. Optimal search for the best alternative. Vol. 78. Department of Energy; 1978.

18. Manso G. Motivating Innovation. J Finance [Internet]. 2011 Oct 1;66(5):1823–60. Available from: 10.1111/j.1540-6261.2011.01688.x

19. Baska M. AI will give workers back two weeks a year, says research. People Management [Internet]. 2018 Sep 14; Available from: https://www.peoplemanagement.co.uk/article/1744734/artificial-intelligence-give-workers-back-two-weeks-year

20. Ortiz-Ospina E, Roser M. Loneliness and social connections. Our World Data. 2023;

21. Bloom N, Jones CI, Van Reenen J, Webb M. Are Ideas Getting Harder to Find? Am Econ Rev [Internet]. 2020;110(4):1104–44. Available from: https://www.aeaweb.org/articles?id=10.1257/aer.20180338

22. Rowell D, Connelly LB. A History of the Term “Moral Hazard.” J Risk Insur [Internet]. 2012 Mar 26;79(4):1051–75. Available from: http://www.jstor.org/stable/23354958

23. Winter RA. Optimal insurance under moral hazard. Handb Insur. 2000;155–83.

24. Chugunova M, Sele D. We and It: An Interdisciplinary Review of the Experimental Evidence on Human-Machine Interaction [Internet]. Vol. 12/2020, Center for Law & Economics Working Paper Series. Center for Law & Economics, ETH Zurich; Available from: http://hdl.handle.net/20.500.11850/442053

25. Schniter E, Shields TW, Sznycer D. Trust in humans and robots: Economically similar but emotionally different. J Econ Psychol [Internet]. 2020;78:102253. Available from: https://www.sciencedirect.com/science/article/pii/S0167487020300106

26. Melo C De, Marsella S, Gratch J. People Do Not Feel Guilty About Exploiting Machines. ACM Trans Comput Interact [Internet]. 2016 May 28;23(2):1–17. Available from: https://dl.acm.org/doi/10.1145/2890495

27. Wickens CD, Clegg BA, Vieane AZ, Sebok AL. Complacency and Automation Bias in the Use of Imperfect Automation. Hum Factors. 2015 Aug;57(5):728–39.

28. Woessner MN, Tacey A, Levinger-Limor A, Parker AG, Levinger I. The evolution of technology and physical inactivity: the good, the bad, and the way forward. Front public Heal. 2021;9:655491.

29. Galletta DF, Durcikova A, Everard A, Jones BM. Does spell-checking software need a warning label? Commun ACM [Internet]. 2005 Jul;48(7):82–6. Available from: https://dl.acm.org/doi/10.1145/1070838.1070841

30. Alberdi E, Povyakalo A, Strigini L, Ayton P. Effects of incorrect computer-aided detection (CAD) output on human decision-making in mammography. Acad Radiol [Internet]. 2004;11(8):909–18. Available from: https://www.sciencedirect.com/science/article/pii/S1076633204003265

31. Lawton-Smith H. Universities, innovation and the economy. Taylor & Francis; 2006.

32. Rudolph J, Tan S, Tan S. ChatGPT: Bullshit spewer or the end of traditional assessments in higher education? J Appl Learn Teach [Internet]. 2023 Jan 25;6(1). Available from: https://journals.sfu.ca/jalt/index.php/jalt/article/view/689

33. Qadir J. Engineering education in the era of ChatGPT: Promise and pitfalls of generative AI for education. In: 2023 IEEE Global Engineering Education Conference (EDUCON). IEEE; 2023. p. 1–9.

34. Tlili A, Shehata B, Adarkwah MA, Bozkurt A, Hickey DT, Huang R, et al. What if the devil is my guardian angel: ChatGPT as a case study of using chatbots in education. Smart Learn Environ [Internet]. 2023;10(1):15. Available from: 10.1186/s40561-023-00237-x

35. Lo CK. What Is the Impact of ChatGPT on Education? A Rapid Review of the Literature. Vol. 13, Education Sciences. 2023.

36. Friederichs H, Friederichs WJ, März M. ChatGPT in medical school: how successful is AI in progress testing? Med Educ Online. 2023 Dec;28(1):2220920.

37. Ederer F, Manso G. Is Pay for Performance Detrimental to Innovation? Manage Sci [Internet]. 2013 Mar 26;59(7):1496–513. Available from: http://www.jstor.org/stable/23443866

38. Crosetto P, Filippin A. The “bomb” risk elicitation task. J Risk Uncertain [Internet]. 2013 Mar 26;47(1):31–65. Available from: http://www.jstor.org/stable/43550175

39. Haas N, Hassan M, Mansour S, Morton RB. Polarizing information and support for reform. J Econ Behav Organ [Internet]. 2021;185:883–901. Available from: https://www.sciencedirect.com/science/article/pii/S0167268120303814

40. Hassan M, Amin E, Mansour S, Voigt S. Incentivizing cooperation against a norm of defection: Experimental Evidence from Egypt. J Behav Exp Econ [Internet]. 2023;107:102121. Available from: https://www.sciencedirect.com/science/article/pii/S2214804323001477

41. Amin E, Abouelela M, Soliman A. The role of heterogeneity and the dynamics of voluntary contributions to public goods: An experimental and agent-based simulation analysis. J Artif Soc Soc Simul. 2018;21(1).

42. Kleinberg J, Lakkaraju H, Leskovec J, Ludwig J, Mullainathan S. Human Decisions and Machine Predictions. Q J Econ [Internet]. 2018 Feb 1;133(1):237–93. Available from: 10.1093/qje/qjx032

43. Hoffman M, Kahn LB, Li D. Discretion in Hiring. Q J Econ [Internet]. 2018 May 1;133(2):765–800. Available from: 10.1093/qje/qjx042

44. Romero-Rodríguez J-M, Ramírez-Montoya M-S, Buenestado-Fernández M, Lara-Lara F. Use of ChatGPT at University as a Tool for Complex Thinking: Students’ Perceived Usefulness. J New Approaches Educ Res Vol 12, No 2 [Internet]. 2023; Available from: https://naerjournal.com/article/view/1458

45. Dyer J, Gregersen H, Christensen C. The Innovator’s DNA: Mastering the Five Skills of Disruptive Innovators. Boston: Harvard Business Review Press; 2011.

